# An evolutionarily conserved role for VEGF signaling in the expansion of non-vascular tissue during regeneration

**DOI:** 10.1101/2024.10.01.616057

**Authors:** Aaron M. Savage, Alexandra C. Wagner, Ryan T. Kim, Paul Gilbert, Hani D. Singer, Erica Chen, Elane M. Kim, Noah Lopez, Kelly E. Dooling, Julia C. Paoli, S.Y. Celeste Wu, Sebastian Bohm, Rachna Chilambi, Tim Froitzheim, Steven J. Blair, Connor Powell, Adnan Abouelela, Anna G. Luong, Kara N. Thornton, Benjamin Tajer, Duygu Payzin-Dogru, Jessica L. Whited

## Abstract

Salamanders are capable of regenerating whole limbs throughout life, a feat that is unmatched within tetrapods. Limb regeneration is dependent upon the formation of a blastema, which contains undifferentiated cells capable of giving rise to most cells of the regenerated limb. Innervation is required for regeneration, along with many signaling pathways, including FGF, BMP and Wnt, but the role of VEGF signaling during salamander limb regeneration is not well understood. Here we show that VEGF signaling is essential for limb regeneration and that blastema cells and limb fibroblasts display impaired proliferation in the absence of VEGF signaling. By performing analogous experiments in planaria, which lack vasculature, we show a potential evolutionarily conserved role for VEGF in the expansion of blastema tissues that is separable from angiogenesis. Moreover, loss of VEGF signaling reduces induction of EMT-like processes, suggesting VEGF signaling functions upstream of the expression of EMT transcription factors, including *Snai2*. These findings highlight potential roles for VEGF signaling during regeneration which may extend beyond typical findings related to angiogenesis.

## Introduction

The axolotl (*Ambystoma mexicanum*), along with other salamanders, is capable of the feat of complete regeneration of functional limbs, including regeneration of all necessary tissue types. For proper limb regeneration to occur, the wound must first be healed and covered by a wound epidermis, which facilitates the coalescence of blastema cells at the site of injury, where they proliferate and eventually differentiate into tissues of the regenerated limb (Mccusker et al., 2015; Tajer et al., 2023). The blastema is a transient structure composed of progenitor cells which give rise to much of a regenerated limb, including bones, muscle, and connective tissues; some regenerated cell types have been shown to form from specific blastema cell lineages (Currie et al., 2016a; Kragl et al., 2009; Leigh et al., 2018). How blastema cells are specified is still not clear. Understanding how some wounds in axolotls heal with simple healing, whereas wounds induced by amputation heal by blastema formation, is critical. This transition may represent a crux to limb regeneration necessary to traverse in future therapeutic applications. Candidate genetic regulatory pathways that might shift a simple-wound-healing response to a blastema-formation response have been proposed from transcriptional analyses (Campbell et al., 2011; Knapp et al., 2013), but few experiments have addressed this possible crux.

Beyond blastema formation, a hallmark of successful regeneration is its scar-free nature. Axolotl limb regeneration leaves no visible trace of injury on the skin, unlike scarring in mammals. However, repeated amputation causes failure to regenerate and differences in wound healing (Bryant et al., 2017a). This shares some characteristics with scarring and my highlight an inability to transition between wound healing and regenerative states. In mammals, wound healing is succeeded by a burst of vascularization, which facilitates fibroblast and immune cell arrival at the wound site, but often results in the formation of a scar (Tonnesen et al., 2000). Moreover, endothelial growth factor (VEGF) signaling is inhibitory to mouse digit tip regeneration (Yu et al., 2014) and VEGF overexpression in dermal fibroblasts results in increased scar formation after transplantation into rats (Shams et al., 2022), suggesting a negative role for VEGF signaling and angiogenesis in this process in mammals. Conversely, in axolotls, VEGF signaling inhibition was shown to be insufficient to impair tail regeneration, while increased VEGF signaling, causing increased angiogenesis, did not block tail regeneration (Brashears et al., 2024; Ritenour and Dickie, 2017). These findings highlight how evolutionarily conserved signaling pathways may be differentially regulated following injury in different species. Here we inhibit VEGF signaling during axolotl limb regeneration and show a loss of limb regeneration but not blastema formation. This we found this VEGF dependency to be conserved in planarian tail regeneration, even though planarians lack vasculature. We found VEGF signaling regulates proliferation in both axolotl blastemas and in cultured axolotl limb fibroblasts. Through sequencing studies, we implicate VEGF-mediated regulation of EMT-like processes in axolotl limb regeneration. These results highlight previously underappreciated roles of VEGF signaling in regeneration beyond its established role during angiogenesis, illustrating the need to further characterize the intricacies of this pathway in regenerative organisms.

## Results

### VEGF signaling is required for limb regeneration

VEGF signaling is most well-characterized in the process of angiogenesis (Astin et al., 2014; Blanco and Gerhardt, 2013; Bower et al., 2017; Cleaver and Krieg, 1998; Hogan and Schulte-Merker, 2017; Koch et al., 2011; Olsson et al., 2006; Phng and Gerhardt, 2009), but little study during limb regeneration has taken place so far. The presence of vasculature has been observed in salamander blastemas but how this impacts blastema growth has not been established (Rageh et al., 2002; Smith and Wolpert, 1975; Whited et al., 2013). A previous study utilized the VEGF receptor inhibitor, PTK787, at 100 nM, and observed no loss of regeneration, despite observing reduced angiogenesis. However, the IC50 of PTK787 for Vegfr1, Vegfr2 and Vefgr3 are 37, 77 and 664 nM, respectively, (Drevs et al., 2002; Ritenour and Dickie, 2017), suggesting that Vefgr3 may compensate for the loss of function of the other two receptors. Furthermore, these results highlight the possibility that the function of VEGF signaling during limb regeneration in salamanders might be more complex than simply regulating angiogenesis and may in fact function during the regrowth of other tissue types.

We therefore sought an alternative inhibitor that could be used at low doses to fully inhibit Vegfr function during axolotl limb regeneration. We used the VEGF receptor tyrosine kinase inhibitor AV951 (Tivozanib) which has been used to block angiogenesis in zebrafish (Kang et al., 2013; Nakamura et al., 2006; Savage et al., 2019). We performed mid-stylopod amputations on axolotls and treated them with 80nM AV951 via immersion from 3 days post amputation (dpa) until 22 dpa, to specifically examine the requirement for angiogenesis post-wound healing, to the proliferative phase of limb regeneration. AV951 inhibits VEGF receptors at a lower IC50 (Vegfr1 = 30 nM, Vegfr2 = 6.5 nM, and Vegfr3 =15 nM), meaning our dosage likely completely blocks Vegfr function. We observed a significant reduction in blastema area in AV951-treated limbs from 11dpa onwards (**Fig. 1A-C**) and a subsequent cessation of blastema progression (**Fig. 1D-G**). We stained limbs with Alcian blue to label cartilage in the regenerate limbs, and we observed no cartilage formation distal to the amputation plane in AV951-treated limbs (**Fig. 1H, I**). This result confirms that VEGF signaling is required following blastema formation and prior to chondrogenesis. To determine if the requirement for VEGF signaling is reversible, we treated limbs until 19 dpa and then removed treatment. Limbs were able to recover and begin regenerating and we measured blastemas until paddle stage, which AV951-treated animals reached by 33 dpa (**Fig. 1J**), indicating that blastemas were able to regenerate as normal, once VEGF signaling resumed.

**Figure 1.**
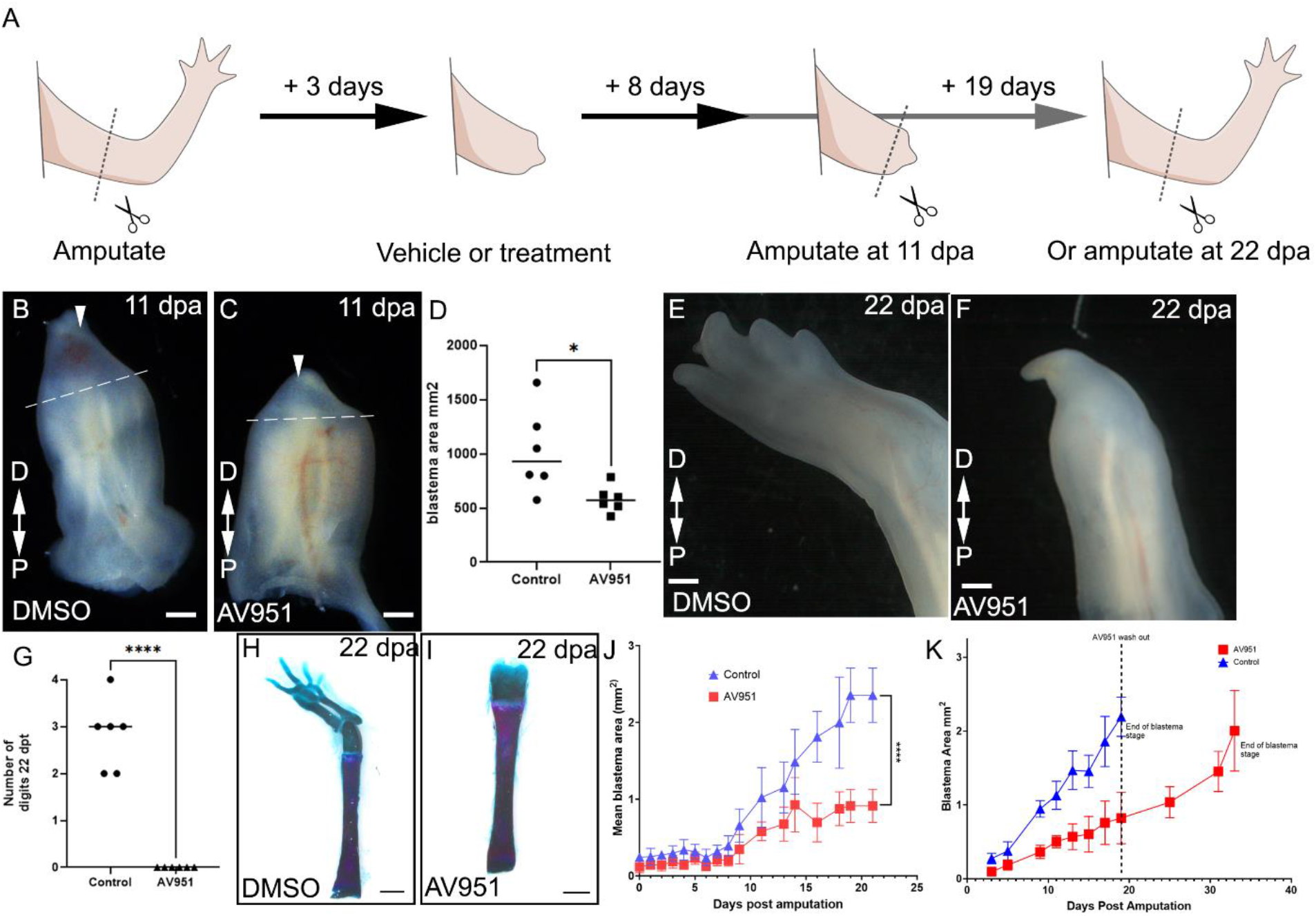
VEGF signaling is required for blastema growth. (A) schematic of experimental design; (B) representative DMSO-treated limb at 11dpa (scale bar 100µm; arrowhead indicates blastema; dashed line indicates amputation plane); (C) representative AV951-treated limb at 11dpa (scale bar 100µm; arrowhead indicates blastema; dashed line indicates amputation plane); (D) AV951-treated blastemas are significantly smaller at 11dpa; (E) representative DMSO-treated limb at 22dpa (scale bar 100µm); (F) representative AV951-treated limb at 22dpa (scale bar 100µm; arrowhead indicates blastema); (G) AV951-treated limbs show no regeneration at 22dpa; (H) skeletal prep of representative DMSO-treated limb at 22dpa (scale bar 250µm); (I) skeletal prep of representative AV95-treated limb at 22dpa (scale bar 250µm); (J) AV951-treated blastemas display blocked regeneration compared to controls; (K) AV951-treated blastemas can develop to paddle stage upon treatment removal. D: distal, P: proximal, dpa: days post-amputation.

VEGF signaling has been primarily associated with angiogenesis during development and disease (Abhinand et al., 2016; Benedito et al., 2012; Bergers and Benjamin, 2003; Hogan and Schulte-Merker, 2017b; Shin et al., 2016a, 2016b) and vasculature has been observed in salamander blastemas (Rageh et al., 2002; Smith and Wolpert, 1975). To determine whether angiogenesis might be impaired following treatment with AV951, we analyzed vascular marker expression in post-amputation limbs. We performed *in situ* hybridization chain reaction (HCR) targeting *Cd34* and measured the total length of contiguous endothelial cells. We observed a significant reduction in *Cd34*^*+*^ vasculature in blastema tissue distal to the amputation plane (**Fig. 2A-C**), suggesting AV951 treatment blocks angiogenesis during axolotls, consistent with findings in zebrafish (Chimote et al., 2014; Savage et al., 2019). Interestingly, despite loss of regeneration, we observed small blastemas in AV951-treated animals; we therefore wanted to determine whether blastema cells were specified at all. We previously identified *Kazald2* (*Kzd2*) as a highly-specific marker of blastema cells (Bryant et al., 2017b; Leigh et al., 2018) and chose this to label presumptive blastemas. Both treatment groups displayed *Kzd2*^*+*^ cells located within the blastema region (**Fig. 2 D-E**). Since a blastema formed but did not grow, we wanted to determine whether proliferation was impaired in AV951-treated blastemas. We therefore administered a short pulse of EdU to label cells in S-phase, and we observed a significant reduction in EdU^+^ nuclei within the blastema of AV951-treated limbs, but no significant difference in EdU^+^ nuclei in tissue more proximal than the amputation plane (**Fig. 2 F-I**). This suggests that blastema cells are capable of coalescing and migrating to the plane of amputation in the absence of VEGF signaling, but they require VEGF signaling to sufficiently proliferate once there.

**Figure 2.**
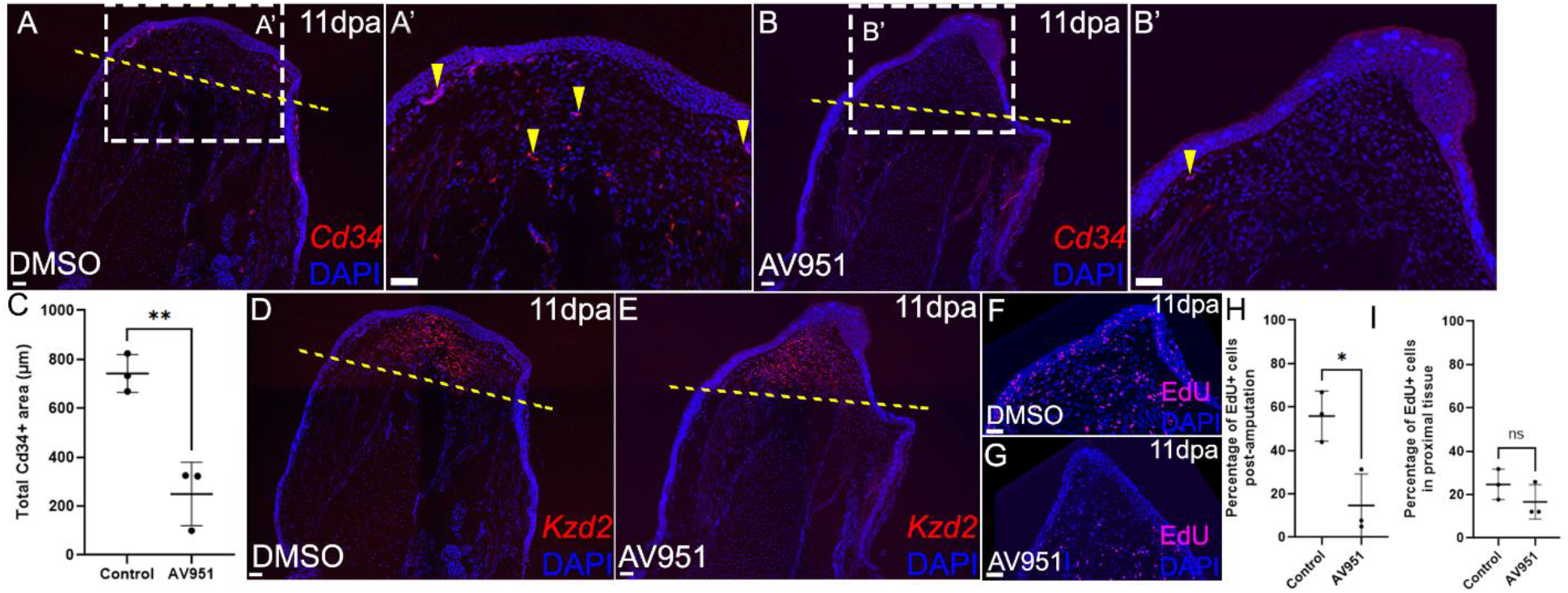
VEGF signaling is required for blastema proliferation but not for blastema cell specification. (A-A’) DMSO-treated blastemas display vascularization beyond the plane of amputation (scale bar 50µm; arrowhead indicates *Cd34*^*+*^ vasculature; dashed line indicates amputation plane); (B-B’) AV951-treated blastemas display little vascularization beyond the plane of amputation (scale bar 50µm; arrowhead indicates *Cd34*^*+*^ vasculature; dashed line indicates amputation plane) (C) AV951-treated blastemas display significantly-reduced vascularization in post-amputation area; (D-E) Blastema cell specification, highlighted by *Kazald2* expression, is not affected by AV951 treatment (scale bar 50µm; dashed line indicates amputation plane); (F-G) AV951-treated blastemas display reduced proliferation; (H-I) reduced proliferation is specifically observed in the distal blastema, not the pre-amputation area.

### VEGF signaling is required in non-vascular tissue during regenerative processes

Our results indicated that blastema cells require VEGF signaling to sufficiently proliferate. One possible explanation of this result is that the requirement is indirect: blastema cells may be dependent upon growth or survival signals supplied by nascent blood vessels, which are also reforming during this time, and vessel regeneration requires VEGF signaling. An alternative, but not mutually-exclusive, possibility, is a direct requirement for VEGF signaling within nascent blastema themselves, many of which are derived from fibroblasts. As the in vivo limb is a complicated composite of many cell types, we tested these possibilities using cultured AL-1 axolotl limb fibroblast cells, which contribute to blastemas upon transplantation into regenerating limbs (Yu et al., 2023), indicating some blastema cell properties are conserved in these cells. We cultured AL-1 cells and performed a scratch assay, wherein cells are removed from a confluent well and regrowth is quantified, which has previously been applied to cultured AL-1 cells (Oliveira et al., 2022). AV951-treated AL-1 cells displayed reduced recovery over the scratch compared to control treated AL1 cells (**Fig. 3 A-I**), suggesting they may directly respond to VEGF signaling. However, whether this is due to migration or proliferation defects—or potentially both—was not clear. We therefore performed an EdU assay to determine whether VEGF signaling could impact AL-1 proliferation. We observed that AL1 cells treated with AV951 displayed impaired proliferation compared to controls (**Fig. 3 J-L**). Our data suggest that AL-1 cells, like blastema cells, also respond to VEGF signaling directly via proliferation, and the lack of wound closure observed in the scratch assay may be due to reduced proliferation, which correlates with our findings *in vivo*.

**Figure 3.**
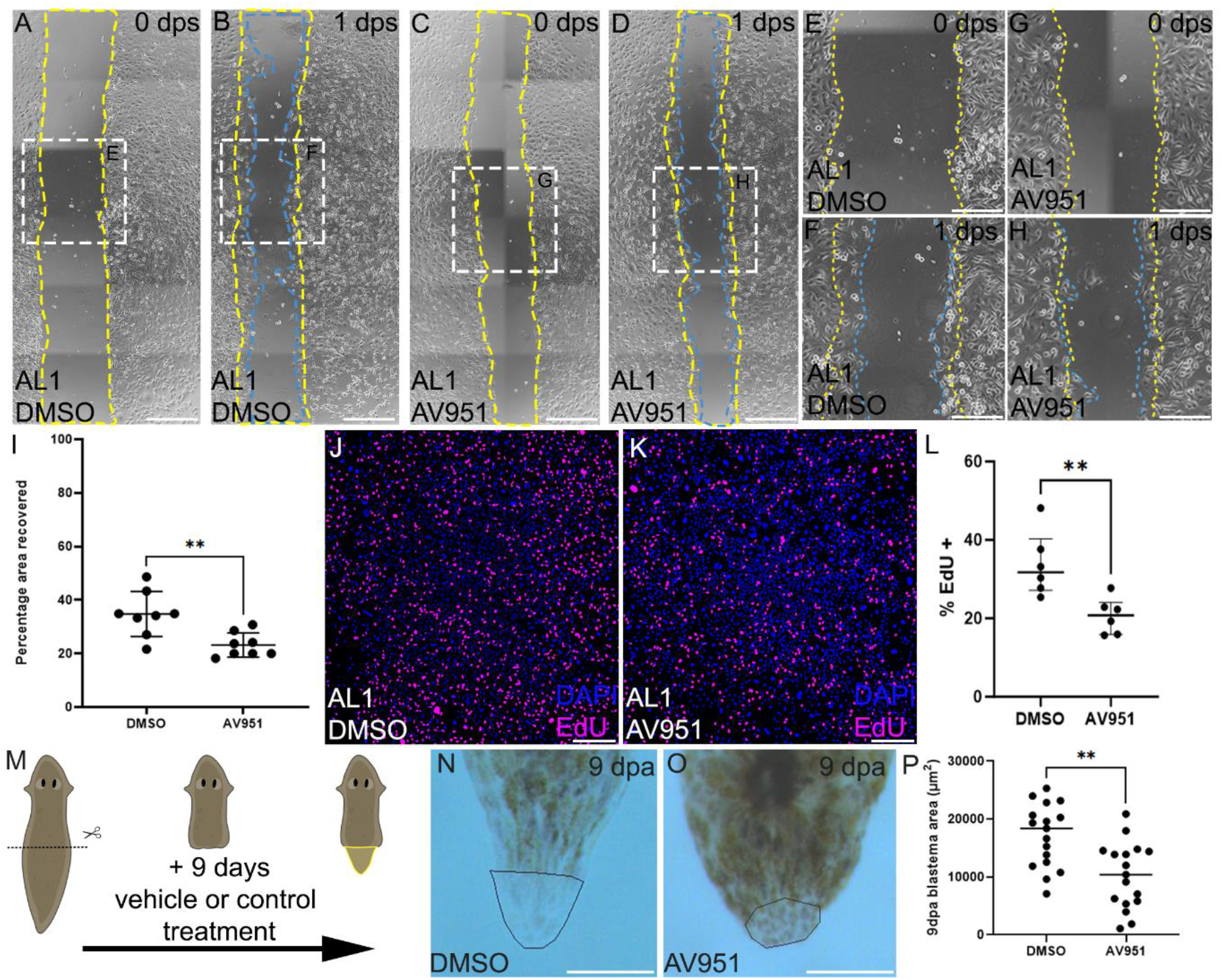
VEGF signaling is required for proliferation of limb fibroblast cells and displays an evolutionarily conserved function. (A) representative DMSO-treated AL1 cells at 0 days post scratch (dps) (yellow dashed line indicates periphery of scratch at 0dps; scale bar 500 µm); (B) representative DMSO-treated AL1 cells at 1dps (yellow dashed line indicates periphery of scratch at 0dps; blue dashed line indicates periphery of scratch at 1 dps; scale bar 500 µm) (C) representative AV951-treated AL1 cells at 0dps (yellow dashed line indicates periphery of scratch at 0dps; scale bar 500 µm); (D) representative AV951-treated AL1 cells at 1dps (yellow dashed line indicates periphery of scratch at 0dps; blue dashed line indicates periphery of scratch at 1 dps scale bar 500 µm); (E-H) inset images from A-D showing closure in different conditions; (I) AV951-treated AL1 cells display a significantly reduced recovery from scratch (scale bar 500 µm); (J) representative image of EdU staining in DMSO-treated AL1 cells (scale bar 500 µm); (K) representative image of EdU staining in AV951-treated AL1 cells (scale bar 500 µm); (L) Av951-teated Al1 cells display significantly reduced proliferation compared to controls; (M) schematic of planaria amputation and treatment; (N-O) representative images of control and AV951-treated planarian blastemas at 9 dpa (scale bar 200µm); (P) quantification of blastema area.

While axolotls represent the most basal group of tetrapod organisms that can regenerate limbs, regeneration is a widespread phenomenon among many animal clades. However, not all regenerating species have vasculature. We hypothesized that VEGF signaling may function convergently in blastema cells of distantly related, highly-regenerative species. We chose planaria because they regenerate large body parts, express a VEGF receptor, termed *Vegfr1*, while lacking a characteristic vascular system (Lei et al., 2016). We treated planaria (*Girardia dorotcephala*) with 80nM AV951 and measured blastema growth at 9dpa (**Fig. 3M**). We observed significantly smaller blastemas in the AV951-treated planarians compared to DMSO-treated control planarians (**Fig. 3N-O**). This result demonstrates that the requirement for VEGF signaling in complex tissue regeneration is conserved across highly-diverged species and that one aspect of this requirement is likely independent of vasculature.

### Loss of VEGF signaling traps blastemas in a wound healing-like state and impairs expression of EMT-like genes

Since our data indicate that VEGF signaling is required for blastema progression, we were interested in determining what the effects of a loss of VEGF signaling were on the blastema at the transcriptional level. We performed Illumina bulk RNA sequencing on blastemas obtained from AV951-treated and control (DMSO-treated) animals and we performed differential gene expression analysis. We identified 3287 differentially expressed genes between the two conditions (**Fig. 4A**), and our PCA analysis showed clustering of the two sample groups (**Fig S1A**), and we were able to observe differential gene expression in a volcano plot (**Fig. S1B**), or heat map (**Fig. 3A**) highlighting downregulation of genes associated with angiogenesis and Wnt signaling, which was corroborated by Kegg analysis (**Fig S2**).

**Figure 4.**
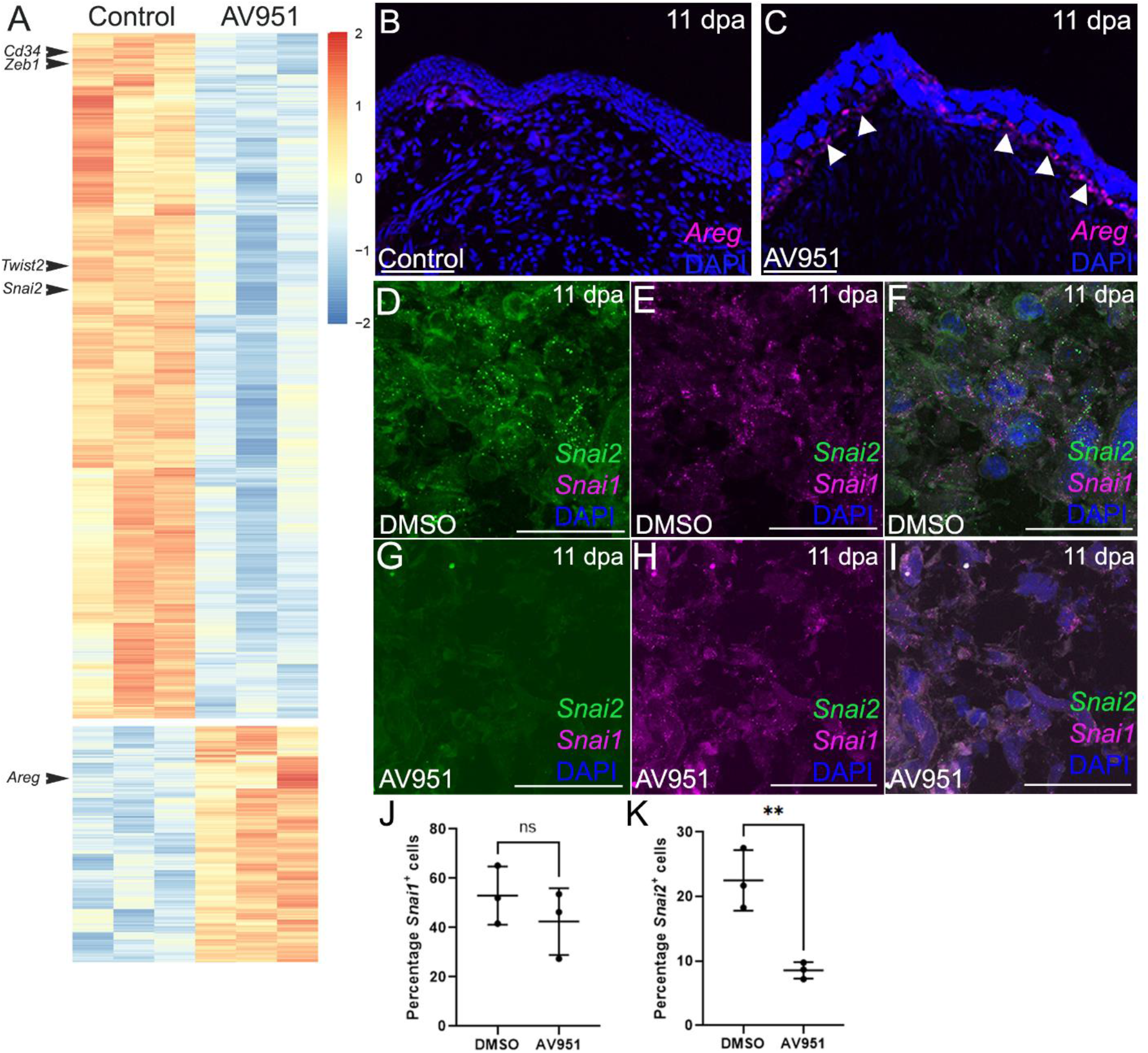
Loss of VEGF signaling induces ectopic expression of early stage wound healing genes and a loss of EMT transcription factor expression. (A) heat map displaying differentially expressed genes in AV951-treated blastemas compared to controls; (B) control treated blastemas display no *Areg* expression at 11 dpa (scale bar 100µm); (C) AV951-treated blastemas display ectopic *Areg* expression at 11 dpa (white arrowheads; scale bar 100µm); (D-F) *Snai1* and *Snai2* expression in mesenchymal cells of control blastema at 60x magnification (scale bar 50µm); (G-I) Av951-treated blastemas display reduced *Snai2* expression in mesenchymal cells of the blastema (scale bar 50µm); (J-K) Av951-treated blastemas display reduced expression of *Snai2*, but not *Snai1*.

We also observed differential expression of genes associated with epidermal differentiation and wound healing, including *Amphiregulin* (*Areg)*. We previously observed ectopic upregulation of *Areg* in limbs subject to repeat amputation which failed to regenerate (Bryant et al., 2017a).We observed ectopic expression of *Areg*, using HCR, in the basal wound epidermis of AV951 treated animals, but not control animals, validating our bulk RNA-seq findings (**Fig. 3B-C**). Since *Areg* expression is transiently observed during wound healing before being extinguished (Bryant et al., 2017a), this suggests that VEGF signaling might be required for blastemas to transition out of the wound healing phase of regeneration.

Interestingly, we also observed significant differential expression of genes associated with epithelial-to-mesenchymal transition (EMT), which has previously been implicated in limb regeneration (Glotzer et al., 2022; Kawakami et al., 2006; Li et al., 2021; Sader et al., 2019; Wischin et al., 2017; Yu et al., 2023).This suggests a loss of proliferative potential induced by loss of VEGF signaling may be, at least in part, caused by a loss of such programs. The role of EMT-like processes during axolotl limb regeneration is not fully clear, but we have recently hypothesized that EMT-like processes are required for the migration of blastema progenitor cells to migrate to the blastema proper during regeneration (Kim and Whited, 2024; Li et al., 2021; Sader et al., 2019). We used HCR to corroborate our transcriptomic data and observed a decrease in the number of cells expressing *Snai2* in AV951-treated limbs, while *Snai1* expression was unchanged (**Fig. 3D-I**), which was consistent with the transcriptomic data (**Fig. 3A**). These findings might suggest that *Snai1* is more important for the initiation of blastema fate, while *Snai2* may regulate maintenance of blastema fate or subsequent proliferation. Given that the mechanisms regarding the activation of EMT-like processes during axolotl limb regeneration are currently unclear, VEGF signaling presents an interesting target for future studies.

## Discussion

Here we present data consistent with a model in which VEGF signaling is required for angiogenesis during axolotl limb regeneration, therefore exerting an indirect effect on the blastema cells that depend on circulatory factors, but that VEGF signaling is also, more directly, required for blastema cell proliferation. Such a dual-role for VEGF signaling in both angiogenesis and fibroblast proliferation has been observed in other biological contexts. Autocrine VEGF signaling from fibroblasts has been observed in some cancers (Goggins et al., 2023; Masood et al., 2001), where it promotes both fibroblast proliferation and angiogenesis. This parallel role for VEGF signaling in a mammalian cancer context is intriguing because tumors have been long-postulated to potentially co-opt molecular pathways that are used constructively by highly-regenerative species to promote blastema formation and growths (Wong and Whited, 2020). Moreover, senescent fibroblasts have been shown to secrete VEGF (Coppe et al., 2006) and a recent publication showed that senescent fibroblasts are required within the blastema to promote regeneration (Yu et al., 2023). This might suggest that senescent fibroblasts are at least in part responsible for maintaining VEGF signaling in the blastema. Moreover, cancer cell lines have been shown to express VEGF receptors and respond to VEGF signaling in an autocrine manner (Abou Faycal et al., 2018; Huang et al., 2019), and cancer-associated fibroblasts (CAFs) have been shown to secrete VEGF to promote angiogenesis (Pape et al., 2020). This might point to a functionally similar role for fibroblast blastema cells, wherein some cells are VEGF responsive, and some secrete VEGF, to regulate both blastema fibroblast growth and the formation of a vascular network required for blastema progression. We also showed that planaria subject to VEGF inhibition display decreased blastema area, consistent with our findings in axolotls. Interestingly, a similar delay in regeneration has been observed in hydra subject to VEGF inhibition (Turwankar and Ghaskadbi, 2019). Since planaria and hydra lack blood vessels, this suggests that the response of blastema cells to VEGF signaling is an ancient, evolutionarily conserved, component of the regenerative program, which warrants further investigation.

We show an increase in expression of genes associated with the early stages of wound healing, such as *Areg*. Ectopic expression of *Areg* causes impaired regeneration (Bryant et al., 2017a). This suggests that some of the pathways associated with the early stages of regeneration may still be active and that blastemas may behave as if wound healing is not complete in the absence of VEGF signaling. Moreover, we observed a decrease in expression of genes associated with epithelial-to-mesenchymal transition (EMT). EMT-related genes have been characterized during limb regeneration in axolotls; *Zeb, Twist*, and *Snail* family gene expression has been observed at different timepoints during regeneration (Sader et al., 2019). Interestingly, our data highlight a downregulation of many genes associated with EMT, including *Zeb1, Zeb2, Twsit2*, and *Snai2*. We also observed a decrease in *Prdx2* expression, which has previously been associated with blastema cells which have taken on an intermediate fate between mesenchymal and epithelial states (Li et al., 2021). Furthermore, cells which have re-entered the cell cycle due to a distant amputation express *Snai1*, while blastema cells and homeostatic cells do not (Payzin-Dogru et al., 2024). We did not observe a difference in *Snai1* expression in our data, but we did observe a difference in *Snai2* expression. These findings might suggest that *Snai1* is more important for the initiation of blastema fate, while *Snai2* may regulate maintenance of blastema fate or subsequent proliferation. Given that the mechanisms regarding the activation of EMT-like processes during axolotl limb regeneration are currently unclear, VEGF signaling presents an interesting target for future studies.

While AV951 has been shown to inhibit angiogenesis, via inhibition of VEGF receptor tyrosine kinase function, it also has potential off-target effects, including the inhibition of *c-Kit* and *Pdgfrb* function (Hepgur et al., 2013), though *c-Kit* inhibition occurs at higher doses, at IC50 78nM in monolayer cell culture. Interestingly, IC50 values for monolayer culture may be inappropriate for three dimensional structures (Berrouet et al., 2020), which suggests higher doses may be required for *in vivo* studies. Inhibition of *Pdgfr* function during axolotl limb regeneration does produce a similar phenotype to inhibition via AV951 (Currie et al., 2016), which means that disentangling this phenotype via genetic manipulation is essential for future studies into the role of VEGF signaling during limb regeneration.

Taken together, our data highlights the effects of loss of VEGF signaling on axolotl limb regeneration. Further study is required, including the use of genetic approaches to block the VEGF signaling response. Overall, our data suggest a model by which the role of VEGF signaling in non-vascular tissues is conserved through evolution, wherein VEGF signaling can regulate the proliferative capacity of non-vascular tissue during the regenerative process. Moreover, VEGF signaling is likely to be required after resolution of wound healing and may function during the regulation of EMT-like processes which facilitate blastema progression. Our work predicts that modulating VEGF signaling post-specification of blastema cells may be a means whereby these important limb progenitor cells can be harnessed for regenerative therapies.

## Materials and methods

### Animal husbandry and surgical procedures

All animal experimentation was approved by and conducted in accordance with Harvard University’s Institutional Animal Care and Use Committee (Protocol # 19-02-346). Age and size-matched leucistic axolotls were used for all animal experiments. Animals were housed in 40% Holtfreter’s Solution at a salinity of 3000 µS, 7.6 pH, and a room temperature of 19°C. For all experiments, animals received bilateral forelimb amputations at the mid stylopod. Animals were anaesthetized in tricaine before amputation and recovered in 19mM sulfamerizine overnight before being returned to 40% Holtfreter’s solution (with or without pharmacological treatment/vehicle) for the remainder of the experiment. Planaria were housed in 0.5 g of instant ocean per 500 mL of deionized water.

### Pharmacological treatments

For blastema AV951 treatments, animals were treated at 80nM AV951 to inhibit the function of VEGFR1, VEGFR2, and VEGFR3 without acute toxicity. AV951 was prepared as a 10mM stock in DMSO and a volume was diluted into 40% Holtfreter’s solution to a concentration of 80nM into which animals were immersed for treatment. An equivalent volume of DMSO was added to 40% Holtfreter’s solution for control treatments. For EdU analysis, animals were subject to a 16-hour EdU pulse. EdU was prepared as a solution of 5µg/µl and diluted to 0.1 µg/µl prior to injection. Animals received 2mg/kg EdU. For AL1 cells, AV951 was used at a concentration of 50nM. EdU treatment was administered at 10mM for 16 hours. For planarians, AV951 was used at a concentration of 80nM in 0.5 g of instant ocean per 500 mL of deionized water.

### RNA extraction

Blastema tissue samples were collected and immediately placed in 500 µl of Trizol reagent (Thermo Fisher, 15596026) in 1.5 ml microcentrifuge tubes. Tissues were homogenized with a handheld drill homogenizer (Pro Scientific, PRO200) for 20 seconds until fully homogenous. 100uL of Chloroform (Sigma, C2432) was added to each sample and mixed thoroughly by hand shaking the tube. After 5 minutes, tubes were spun at 12,000 rpm at 4°C in a microcentrifuge to promote phase separation. Aqueous phase was collected carefully and placed in a new 1.5 ml microcentrifuge tube, then equal volume of 100% Ethanol was added. The mixture was then added to columns from the RNA Clean & Concentrator Kit (Zymo, R1013), and the subsequent cleanup steps from the protocol were followed. Samples were eluted in 15 ul of RNAse-free water, and measured on a Nanodrop One (Thermo Fisher, ND-ONE-W) to confirm A260/A280 above 2.0 and A260/A230 values were above 1.8. Extracted total RNA was measured for quality on an Agilent Tapestation 4200 (Agilent, G2991BA) to confirm RIN scores were above 9.0. Qubit RNA BR Assay Kit (Invitrogen, Q10210) was used to measure RNA concentration on a Qubit 4 Fluoremeter (Thermo Fisher, Q33238). Samples were normalized to 5ng/ul in 60 ul and submitted to the Bauer Sequencing Core Facility at Harvard University.

### RNAseq analysis

Library preparation was done using the Kapa mRNA HyperPrep kit and sequenced on an Illumina NextSeq 1000 at 50 bp read length through the Bauer Sequencing Core Facility at Harvard University. Fastq files were received from the Core and paired-end sequencing reads were quality-trimmed using Trimmomatic (v0.39). Kallisto (v22.04) was used to align reads to the AmexT_v47 transcriptome assembly (Nowoshilow et al., 2018). To find the differentially expressed genes between DMSO and AV951 conditions, *DESeq2* was used (Love et. al, 2014) as a package in RStudio (v2023.03.0).

### Histology

Tissues were fixed in 4% paraformaldehyde in PBS, dehydrated into methanol in series, and stored at −20 °C. Upon use, samples rehydrated and were left overnight in 30% sucrose in PBS. Samples were embedded in optimal cutting temperature compound (O.C.T.) and stored at −80 °C. Samples were sectioned at a thickness of 16 µm.

### In situ hybridization chain reaction

*In situ* hybridization chain reaction (HCR) was performed as previously described (Lovely et al., 2022). Briefly, sectioned blastemas were washed in 2X SSC, treated with tissue clearing solution, washed in 2X SSC, incubated at 37°C in probe hybridization buffer (Molecular Instruments) before incubating overnight at 37°C in probe solution (1:200 dilution of probe in probe hybridization buffer). The next day, slides are washed in probe wash solution (Molecular Instruments) before washing in 5X SSCT and incubating at room temperature in amplification buffer (Molecular Instruments) and incubating overnight at room temperature in hairpin solution (2µl of each hairpin in amplification buffer). Slides are then washed in 5X SSCT, stained with DAPI (1:1000 in PBS), and mounted for imaging.

## Author Contributions

Conceptualization: A.M.S., J.L.W.; Methodology: A.M.S., A.C.W., R.T.K., H.D.S., A.A., S.J.B.; Validation: A.M.S., A.C.W., R.T.K., E.M.K., E.C., H.D.S., A.A.; Investigation: A.M.S., A.C.W., R.T.K., E.M.K., E.C., H.D.S., A.A. P.G., N.L., K.E.D., J.C.P., S.Y.C.W., S.B., R.C., T.F., C.P., A.G.L.; Data curation: K.E.D., R.T.K., E.M.K., E.C., J.P., A.G.L., S.Y.C.W., H.S. B.T., K.N.T; Writing, review & editing: A.M.S., A.C.W., R.T.K., K.E.D., J.C.P., S.Y.C.W., D.P., B.T., J.L.W.; Funding acquisition: A.M.S., J.L.W.

## Acknowledgements

We would like to express our gratitude to Isaac Adatto, Damian Bernard, Brianna Blackmore, Nicholas Cardelia, Hayden Graham, Lauryn Wilson, Omenma Abengowe, Erin Anderson, Rui Qun Miao, and Vicky Yan for their assistance with animal care. We are grateful to members of the Whited Lab for their valuable advice and discussions during this study.

## Supplementary Information

### Supplementary Figures

**Figure S1.**
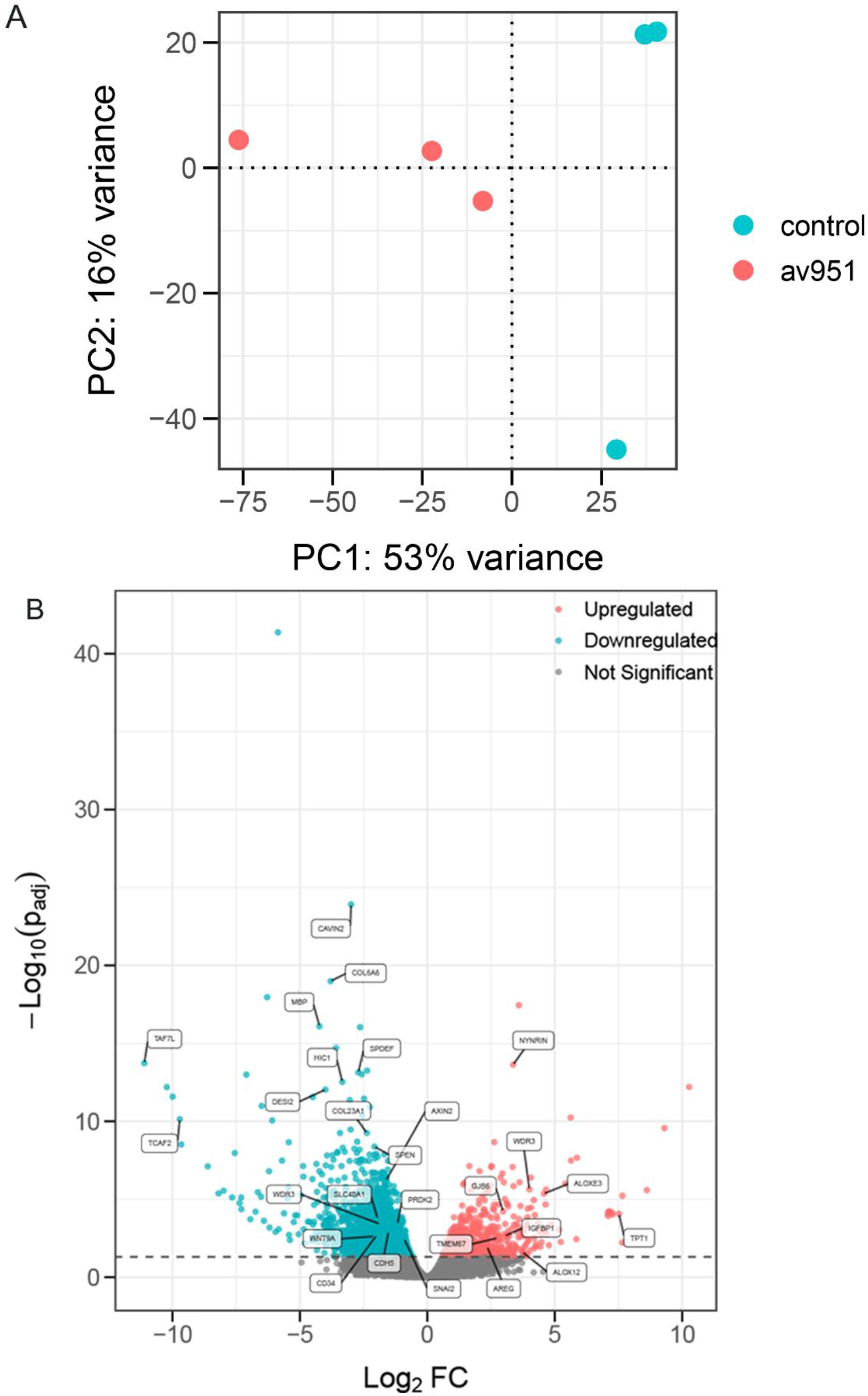
AV951-treated blastemas display significantly differently expressed genes associated with cancer and angiogenesis.

**Figure S2.**
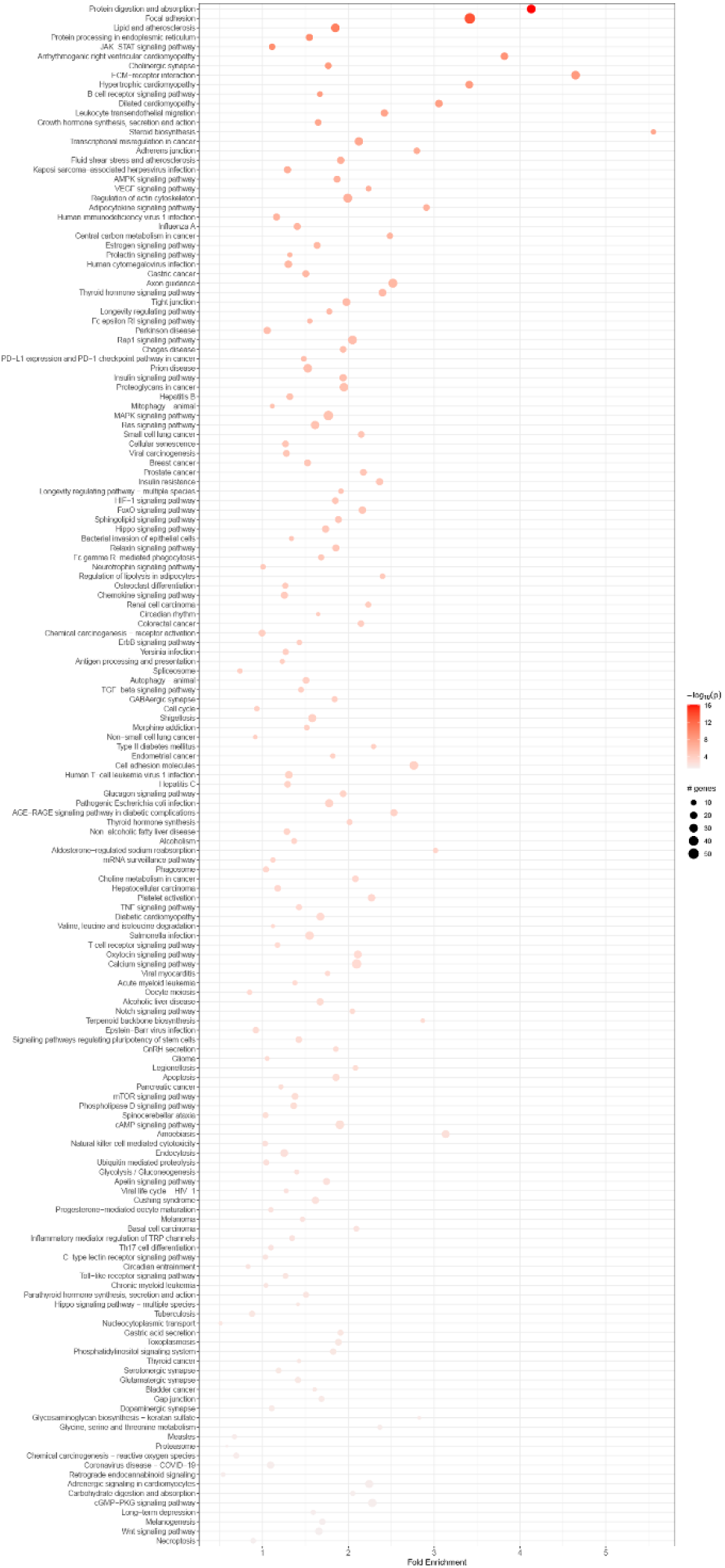
KEGG analysis of bulk RNA seq data highlights developmental pathways which are differentially regulated in AV951-treated blastemas.

